# Understanding detection probability in a low-density population to inform optimal eDNA sampling for freshwater mussels

**DOI:** 10.1101/2025.01.06.631567

**Authors:** Torrey W. Rodgers, Karen E. Mock

## Abstract

Environmental DNA (eDNA) offers a sensitive tool for detecting aquatic species, including those at low population densities. However, effective use of eDNA in conservation and monitoring efforts requires an understanding of detection probabilities and optimal replication effort necessary to minimize false negatives. This study assessed the probability of detecting *Anodonta nuttalliana*, a native freshwater mussel, in two Utah populations with contrasting densities (medium and low). We conducted traditional visual sampling alongside highly replicated eDNA sampling at both sites. In the medium-density population, eDNA provided high detection probabilities with minimal replication. However, in the low-density population, substantially greater replication was needed to achieve similar detection probabilities. Despite the increased eDNA replication effort needed to detect the low-density population, eDNA outperformed traditional visual sampling, which failed to detect live mussels at the low-density site. Our findings highlight the critical balance between sampling effort and detection success, offering guidance for optimizing eDNA sampling strategies in monitoring efforts, especially for species which occur at low densities.

## Introduction

Understanding detection probability is crucial for optimizing sampling intensity in environmental DNA (eDNA) studies. Without accounting for detection probability, researchers risk underestimating species distributions due to false negatives, which can bias ecological assessments and species monitoring when a species is not detected even when it is present at a site. This is especially important in low density populations, which are often targeted by eDNA studies due to the sensitivity of eDNA methods. In low density populations, however, greater replication may be needed to achieve reasonable detection probability and minimize false negatives. However, replication can be costly, so researchers must strike a balance between cost and detection probability.

To this end, we conducted replicated, fine scale eDNA sampling, alongside traditional sampling, in two populations of the native freshwater mussel species *Anodonta nuttalliana* (synonym *Anodonta Californiensis;* Chong et al. 2008) that vary in population density. We sampled both a medium-density population, and a low-density population, to examine and compare detecting probabilities at these two sites, with the goal informing how much replication is needed to detect such populations with high probability.

Environmental DNA sampling is a reliable method for detecting native freshwater mussels from water samples (Stoeckle et al. 2016, Dysthe et al. 2018, Rodgers et al. 2020, 2022). This tool is valuable for determining distributions, and for monitoring of these species, many of which are threatened or endangered. Additionally, native freshwater mussels are often used as indicator taxa in water quality assessments, and their presence or absence can have implications for water quality regulation (Augspurger et al. 2003). eDNA methods are highly sensitive and can detect just a few copies of mitochondrial DNA from a sample. Thus, for high density mussel populations, eDNA can provide high probabilities of detection. For low density populations, however, eDNA may be diluted such that greater sample or technical replication may be necessary to detect mussels with high probability. For species that are rare or threatened, most populations may be at low densities and thus more challenging to detect. Concurrently, low-density populations may be the most important to detect for conservation or regulatory purposes. Thus, knowledge of the amount of replication necessary to detect low density populations with eDNA while maintaining low false negative rates is of importance for the design and implementation of any eDNA monitoring program.

## Methods

### Study sites

eDNA samples were collected from two streams in Utah, the Raft River in NE Utah, and Currant Creek in Central Utah (Figure 1). The Raft River is a tributary of the Snake River and has a medium density of *A. nuttalliana* (high density relative to other populations in Utah). Currant Creek flows into Utah lake and possesses a very low density of *A. nuttalliana*. Although both populations are below agricultural areas, Currant Creek is highly degraded, and impacted by agricultural runoff, cattle grazing, and high densities of invasive Asian clams *Corbicula fluminea* (Richards 2017).Two live *A. nuttalliana* specimens were documented in Currant Creek in 2016 using a clam rake adjacent to where we collected eDNA samples, the year previous to our sampling (Richards 2017).

**Figure 1.**
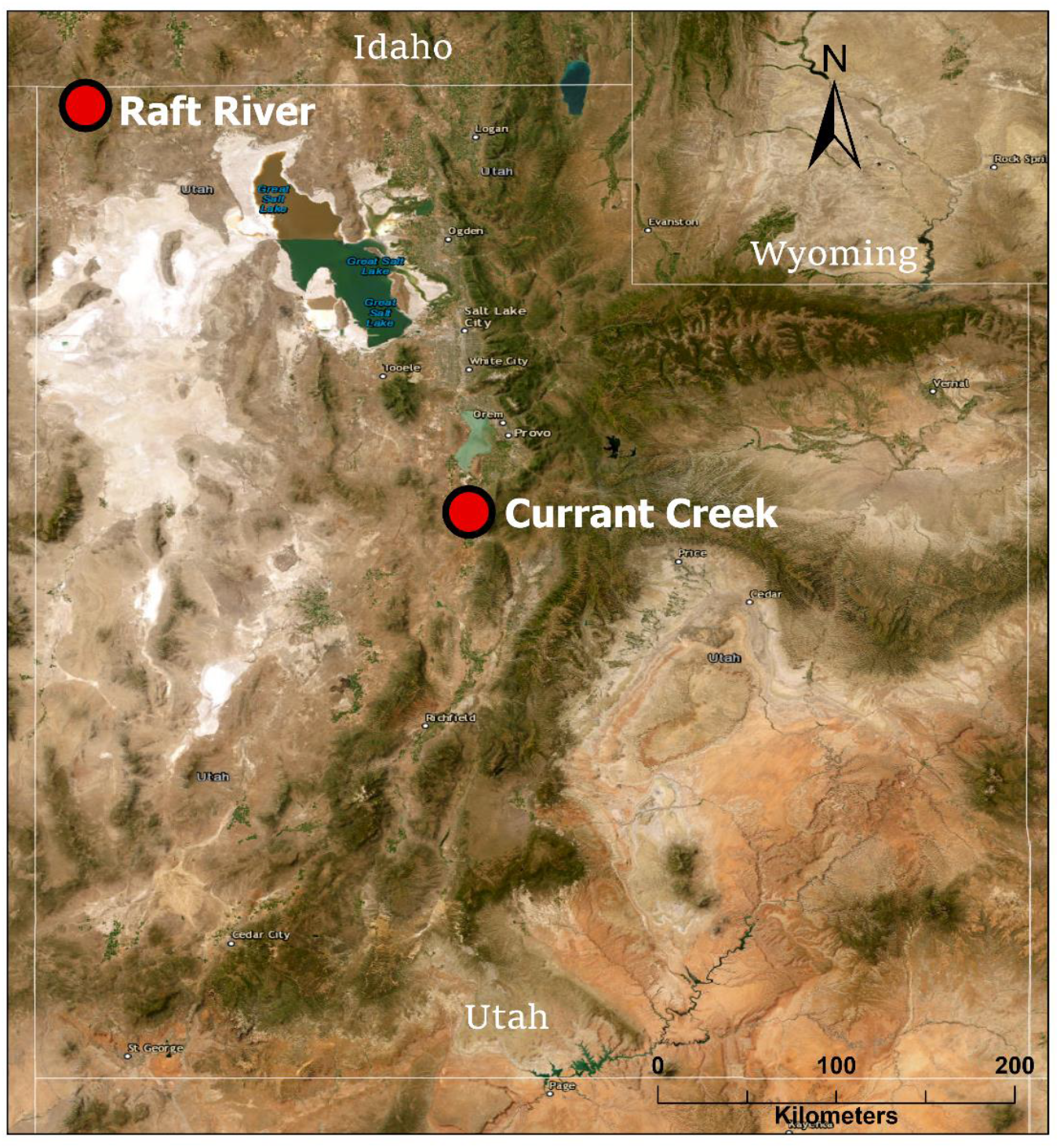
Locations of eDNA and traditional sampling for *Anodonta nuttalliana* in Utah.

### Traditional sampling

At both sites, we conducted traditional visual sampling along a 1 km stretch of stream. The stretch was divided into 50-meter reaches, and each 50 m reach was searched for 15-20 minutes with a glass bottom bucket for freshwater mussels and shells. The number of live individuals and shells were recorded for each site, and we measured the length of all live mussels.

### eDNA sampling

At both sites, an eDNA sample was collected at the downstream end of each 50 m reach for a total of 20 eDNA samples. All samples were collected following the protocol outlined in (Carim et al. 2016). Water was pumped through a 1.5 µm glass microfiber filter using a peristaltic pump until the filter clogged, and filters were stored in silica desiccant until laboratory processing. The volume of water filtered for each sample was recorded. Samples were extracted in a room dedicated for this purpose using the DNeasy Blood & Tissue Kit (Qiagen, Inc. Valencia, CA, USA) with a modified protocol described in Rodgers et al. (2023). Each round of eDNA extraction included one ‘extraction blank’ negative control consisting of an unused sterile filter. All samples were analyzed in nine qPCR replicates following the TaqMan qPCR assay and conditions described in Rodgers et al. (2020). Each qPCR run included six ‘no template’ negative control reactions to monitor for cross-contamination. eDNA extractions were conducted in a dedicated clean lab space physically separated from post-PCR spaces and tissue DNA extraction spaces, and all qPCR reactions were set up under a dedicated PCR hood sterilized with UV radiation prior to each qPCR run (Goldberg et al. 2016).

### Statistical analysis

To examine detection probability at each level of replication, we calculated cumulative binomial detection probabilities for each level of replication across the 20 replicates from each site, and the nine qPCR replicates from each sample (Figure 2c and 2d). To calculate cumulative binomial detection probabilities, we used the following formula:

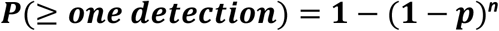

**Figure 2.**
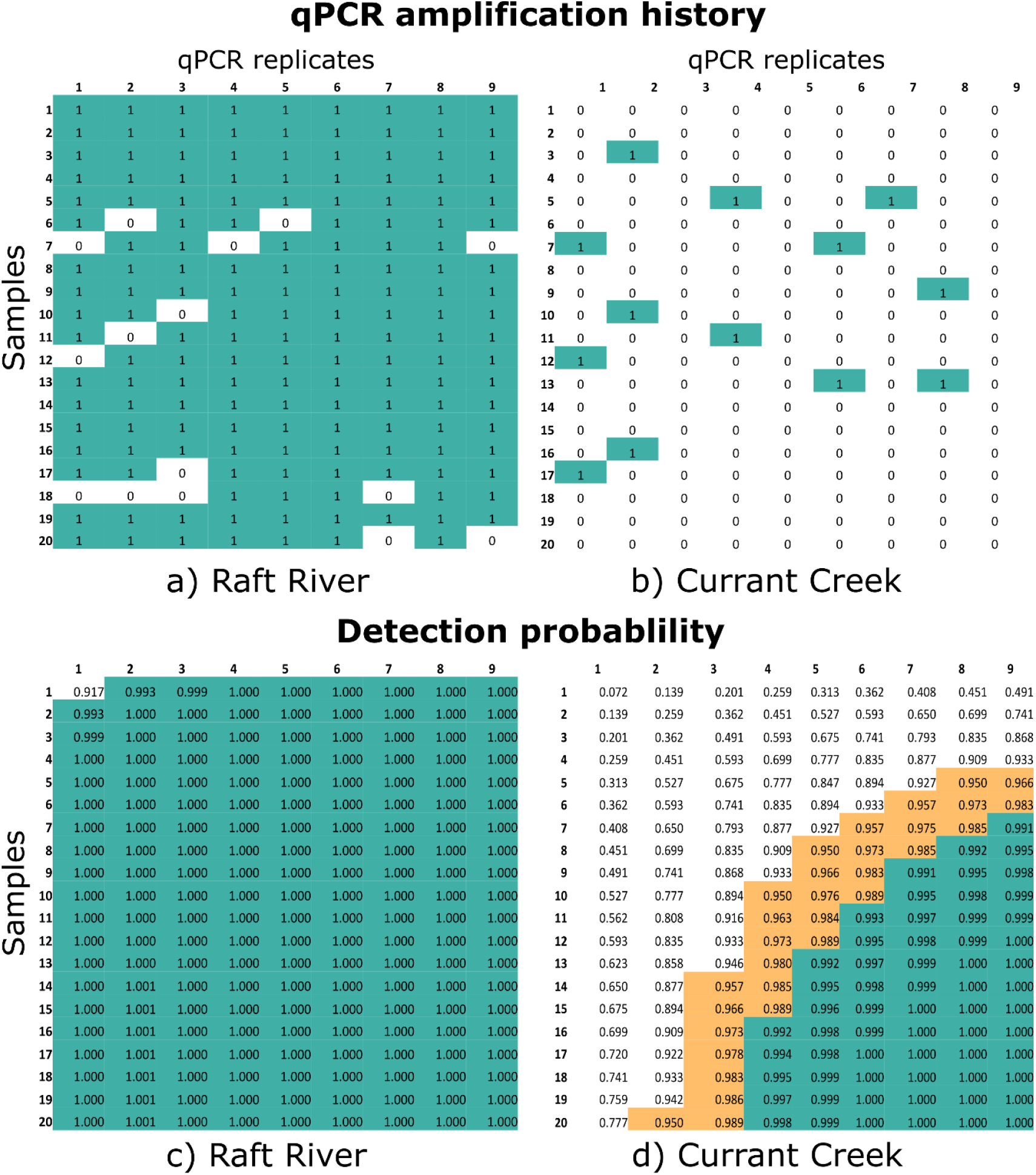
a) Quantitative PCR amplification history for 20 eDNA samples run with nine qPCR replicates for detection of *Anodonta nuttalliana* in the Raft River, Box Elder County, UT, and b) in Currant Creek, Juab County, UT. Turquoise cells represent amplification, and cells in white represent no amplification. c) Binomial detection probabilities for 20 eDNA samples run with nine qPCR replicates in the Raft River, and d) in Currant Creek. Tan cells represent binomial detection probabilities of >0.95, and turquoise cells represent probabilities >0.99.

Where ***P*** is the probability of at least one detection, ***p***is the probability of detection on a given trial (in our case the probability of any given qPCR replicate amplifying), and ***n*** is the number of trials (in our case the number of eDNA samples and/or qPCR replicates).

## Results

### Traditional sampling

In the Raft River, traditional sampling encountered 34 live mussels and 11 shells. Live mussels ranged in maximum length from 11-55 mm (Supplementary table S1). Live mussels were encountered in 13/21 of the 50 m reaches sampled, with 1-6 live mussels detected per reach. In Currant Creek, no live mussels were found with traditional sampling. Thirteen *Anodonta* shells and shell fragments were encountered, including one fully intact and hinged shell (Supplementary table S1).

### eDNA sampling

In the Raft River, Environmental DNA was detected in 20/20 samples, with eDNA concentration estimates ranging from 7.73 - 551.83 DNA copies per liter (Supplementary table S2). All 9 qPCR replicates amplified in 12/20 samples from the Raft River, in four samples, 8/9 replicates amplified, in 3 samples 7/9 replicates amplified, and in one sample, 5/9 replicates amplified (Figure 2a). Overall, from the Raft River, 91.7% of all qPCR replicates amplified (165/180).

In Currant Creek eDNA was detected in 10/20 samples, with concentration estimates ranging from 2.02 – 62.08 DNA copies per Liter (Supplementary table S2). However, in Currant Creek, from the 10 samples where mussel eDNA was detected, in seven samples, just 1/9 qPCR replicates amplified, and in three samples, 2/9 qPCR replicates amplified. From Currant Creek, no more than two of the nine qPCR replicates amplified (Figure 2b) from any of the 20 samples. Overall, from Currant Creek, 7.2% of all qPCR replicates amplified (13/180).

When amplification histories were converted to cumulative binomial detection probabilities to examine detection probabilities across each level of replication, from the Raft River, cumulative binomial detection probabilities ranged from 0.917 (one sample with one qPCR replicate) to 1.0 (from two samples with two qPCR replicates, all the way to 20 samples at 9 qPCR replicates; Figure 2c). From Currant Creek, cumulative binomial detection probabilities ranged from 0.0712 (one sample with one qPCR replicate) to 1.0 (from 15 samples with seven qPCR replicates, to 20 samples at 9 qPCR replicates; Figure 2d). From the Raft River, just two qPCR replicates from one sample, or one qPCR replicate from two samples was needed to achieve a cumulative binomial detection probability of 0.99 (Figure 2c). Conversely in Currant Creek, to achieve a cumulative binomial detection probability of 0.95, two qPCR replicates from all 20 samples, or eight qPCR replicates from five samples were needed (Figure 2d). To achieve a cumulative binomial detection probability of 0.99 in the Currant Creek population, four qPCR replicates from all 20 samples, or all 9 qPCR replicates from seven samples were needed (Figure 2d).

## Discussion

The results of this study highlight the critical importance of sampling replication when using eDNA to detect low-density populations of species like *Anodonta nuttalliana*. Consistent with previous studies, our findings confirm that eDNA is an effective tool for detecting freshwater mussels (Stoeckle et al. 2016, Dysthe et al. 2018, Mauvisseau et al. 2019, Rodgers et al. 2020, 2022), particularly in higher-density populations such as those found in the Raft River. However, in lower-density populations, like Currant Creek, achieving high detection probability requires substantially greater replication.

The stark contrast in detection probabilities between the Raft River and Currant Creek demonstrates how population density influences the success of eDNA detection. In the medium-density Raft River population, high detection probabilities were achieved with low levels of replication. Even with minimal qPCR replication, cumulative binomial detection probabilities approached or exceeded 0.99. In contrast, the low-density population in Currant Creek required much higher levels of replication to achieve similar detection probabilities. This result reflects the lower concentrations of DNA in low-density populations, where the DNA is likely more diluted and harder to detect. These findings are consistent with previous eDNA research indicating that detection probability is reduced in low-density populations, and dependent on both sample and technical replication (Dejean et al. 2012, Schmidt et al. 2013, Furlan et al. 2016, Piggott 2016, Dougherty et al. 2016)

Even though considerable replication was needed to detect *A. nuttalliana* with eDNA in Currant Creek, eDNA sampling still outperformed traditional visual sampling, which failed to detect any live *A. nuttalliana* specimens in five hours of searching. Although no live specimens were found by our traditional sampling, Richards (2017) found 2 live *A. nuttalliana* specimens in Currant Creek in 2016 using a clam rake adjacent to where we collected eDNA samples, the year previous to our sampling. Traditional sampling in Currant Creek was challenging, as the water was cloudy and had poor visibility, the gradient was steep and overgrown in some sections, and the substrate was dominated by shells and live Asian clams *(Corbicula fluminea)*. Thus, Currant Creek epitomizes a situation where eDNA sampling is advantageous for detecting a low-density population, even if a considerable amount of replication was needed to achieve detection, as the population would have been missed completely by traditional sampling, even with five hours of effort.

Our results emphasize the need to carefully balance the trade-offs between cost and detection probability. In Currant Creek, achieving a cumulative binomial detection probability of 0.99 required either four qPCR replicates from all 20 samples or all nine replicates from seven samples. Both scenarios involve considerable laboratory and/or field replication, highlighting the increased effort associated with detecting low-density populations. Researchers and conservation practitioners must weigh these costs against the potential consequences of failing to detect rare or threatened species. In cases where detecting low-density populations is critical for conservation or management decisions, increased replication may be justified despite the higher associated costs.

In conclusion, this study provides valuable insights into optimizing eDNA sampling strategies for detecting freshwater mussels, as well as other aquatic species, particularly in low-density populations. The findings underscore the need for increased replication when density is expected to be low, such as is often the case for threatened or endangered species, as well as recently introduced invasive species. Future research could explore the cost-effectiveness of various replication strategies across different species and habitats to further inform conservation and monitoring practices.

## Acknowledgements

We would like to thank Krissy Wilson from the Utah Division of Wildlife Resources for seeing the value of using eDNA for western freshwater mussels, and for helping fund this study. We acknowledge the use of AI-assisted tools in the editing and writing of some portions of this manuscript.

## Notes

### Competing Interest Statement

The authors have declared no competing interest.

