## Supplementary table S1 for "Understanding detection probability in a low-density population to inform optimal eDNA sampling for freshwater mussels"

**Supplementary Information**

**Table S1**. Results from traditional sampling for *Anodonta nuttalliana* from 20 50-meter reaches from the Raft River in Box Elder County, and 20 50-meter reaches from Currant Creek in Juab County, UT.

| **Sample** | **Water body** | **# Unhinged shells** | **# Hinged shells** | **# Live mussels** | **Live mussel sizes (mm)** |
| --- | --- | --- | --- | --- | --- |
| **RaftFS-01** | Raft River | 0 | 0 | 0 |  |
| **RaftFS-02** | Raft River | 0 | 0 | 0 |  |
| **RaftFS-03** | Raft River | 0 | 0 | 0 |  |
| **RaftFS-04** | Raft River | 1 | 1 | 0 |  |
| **RaftFS-05** | Raft River | 0 | 1 | 1 | 36 |
| **RaftFS-06** | Raft River | 2 | 0 | 5 | 30, 38, 45, 48, 55 |
| **RaftFS-07** | Raft River | 0 | 0 | 0 |  |
| **RaftFS-08** | Raft River | 0 | 0 | 2 | 22, 45 |
| **RaftFS-09** | Raft River | 0 | 0 | 2 | 23, 24 |
| **RaftFS-10** | Raft River | 0 | 0 | 6 | 38, 41, 44, 45, 48, 50 |
| **RaftFS-11** | Raft River | 0 | 2 | 2 | 40, 48 |
| **RaftFS-12** | Raft River | 0 | 1 | 0 |  |
| **RaftFS-13** | Raft River | 0 | 0 | 1 | 42 |
| **RaftFS-14** | Raft River | 0 | 0 | 1 | 40 |
| **RaftFS-15** | Raft River | 0 | 0 | 2 | 20, 25 |
| **RaftFS-16** | Raft River | 0 | 0 | 3 | 20, 27, 39, |
| **RaftFS-17** | Raft River | 0 | 0 | 0 |  |
| **RaftFS-18** | Raft River | 0 | 0 | 1 | 20 |
| **RaftFS-19** | Raft River | 1 | 2 | 6 | 11, 22, 42, 43, 44, 50 |
| **RaftFS-20** | Raft River | 0 | 0 | 2 | 47, 47 |
| **RaftFS-21** | Raft River | 0 | 0 | 1 | 42 |
| **Raft River** | **Totals** | **4** | **7** | **35** |  |
| **CurFS-01** | Currant Creek | 0 | 0 | 0 |  |
| **CurFS-02** | Currant Creek | 0 | 0 | 0 |  |
| **CurFS-03** | Currant Creek | 0 | 0 | 0 |  |
| **CurFS-04** | Currant Creek | 0 | 0 | 0 |  |
| **CurFS-05** | Currant Creek | 0 | 0 | 0 |  |
| **CurFS-06** | Currant Creek | 0 | 0 | 0 |  |
| **CurFS-07** | Currant Creek | 0 | 0 | 0 |  |
| **CurFS-08** | Currant Creek | 0 | 0 | 0 |  |
| **CurFS-09** | Currant Creek | 0 | 0 | 0 |  |
| **CurFS-10** | Currant Creek | 0 | 0 | 0 |  |
| **CurFS-11** | Currant Creek | 0 | 0 | 0 |  |
| **CurFS-12** | Currant Creek | 0 | 0 | 0 |  |
| **CurFS-13** | Currant Creek | 0 | 0 | 0 |  |
| **CurFS-14** | Currant Creek | 1 | 0 | 0 |  |
| **CurFS-15** | Currant Creek | 1 | 0 | 0 |  |
| **CurFS-16** | Currant Creek | 3 | 1 | 0 |  |
| **CurFS-17** | Currant Creek | 1 | 0 | 0 |  |
| **CurFS-18** | Currant Creek | 1 | 0 | 0 |  |

**Table S2.** Results from fine scale environmental DNA sampling for *Anodonta nuttalliana* from the Raft River in Box Elder County, and Currant Creek in Juab County, UT. Samples were collected 50 meters apart.

| **Sample** | **Water body** | **Latitude** | **Longitude** | **Volume (L)** | **Mean Ct** | **Replicates Amplified** | **Estimated Copies/L** |
| --- | --- | --- | --- | --- | --- | --- | --- |
| **RaftFS-01** | Raft River | 41.9632 | -113.6629 | 1.3 | 36.96 | 9 | 170.55 |
| **RaftFS-02** | Raft River | 41.9512 | -113.6694 | 1.7 | 37.45 | 9 | 143.15 |
| **RaftFS-03** | Raft River | 41.9509 | -113.6698 | 2.6 | 36.65 | 9 | 143.70 |
| **RaftFS-04** | Raft River | 41.9505 | -113.6702 | 2.8 | 35.44 | 9 | 251.07 |
| **RaftFS-05** | Raft River | 41.9508 | -113.6707 | 2.2 | 36.53 | 9 | 167.84 |
| **RaftFS-06** | Raft River | 41.9512 | -113.6711 | 3 | 38.17 | 7 | 28.03 |
| **RaftFS-07** | Raft River | 41.9515 | -113.6716 | 2 | 38.96 | 6 | 22.20 |
| **RaftFS-08** | Raft River | 41.9520 | -113.6717 | 2 | 37.47 | 9 | 72.58 |
| **RaftFS-09** | Raft River | 41.9519 | -113.6723 | 1.5 | 38.25 | 9 | 51.75 |
| **RaftFS-10** | Raft River | 41.9523 | -113.6724 | 2.6 | 36.60 | 8 | 61.61 |
| **RaftFS-11** | Raft River | 41.9526 | -113.6724 | 2.5 | 38.00 | 8 | 24.77 |
| **RaftFS-12** | Raft River | 41.9526 | -113.6729 | 2 | 38.65 | 8 | 20.57 |
| **RaftFS-13** | Raft River | 41.9529 | -113.6731 | 2.9 | 37.26 | 9 | 52.64 |
| **RaftFS-14** | Raft River | 41.9530 | -113.6736 | 2.1 | 38.81 | 9 | 35.15 |
| **RaftFS-15** | Raft River | 41.9530 | -113.6741 | 2.7 | 39.16 | 9 | 16.53 |
| **RaftFS-16** | Raft River | 41.9528 | -113.6745 | 0.8 | 36.00 | 9 | 551.83 |
| **RaftFS-17** | Raft River | 41.9524 | -113.6748 | 2.2 | 40.02 | 8 | 7.73 |
| **RaftFS-18** | Raft River | 41.9526 | -113.6751 | 2.1 | 0.00 | 5 | 0.00 |
| **RaftFS-19** | Raft River | 41.9523 | -113.6753 | 2 | 37.21 | 9 | 90.99 |
| **RaftFS-20** | Raft River | 41.9523 | -113.6759 | 2.2 | 38.11 | 7 | 49.94 |
| **CurFS-01** | Currant Creek | 39.9014 | -111.8934 | 0.5 | 0.00 | 0 | 0.00 |
| **CurFS-02** | Currant Creek | 39.9012 | -111.8929 | 0.5 | 0.00 | 0 | 0.00 |
| **CurFS-03** | Currant Creek | 39.9008 | -111.8925 | 0.5 | 39.12 | 1 | 29.73 |
| **CurFS-04** | Currant Creek | 39.9005 | -111.8920 | 0.5 | 0.00 | 0 | 0.00 |
| **CurFS-05** | Currant Creek | 39.9002 | -111.8918 | 0.5 | 0.00 | 2 | 0.00 |
| **CurFS-06** | Currant Creek | 39.8999 | -111.8914 | 0.5 | 0.00 | 0 | 0.00 |
| **CurFS-07** | Currant Creek | 39.8997 | -111.8909 | 0.5 | 37.95 | 2 | 62.08 |
| **CurFS-08** | Currant Creek | 39.8997 | -111.8903 | 0.5 | 0.00 | 0 | 0.00 |
| **CurFS-09** | Currant Creek | 39.8995 | -111.8898 | 0.5 | 0.00 | 1 | 0.00 |
| **CurFS-10** | Currant Creek | 39.8991 | -111.8894 | 0.5 | 38.02 | 1 | 59.51 |
| **CurFS-11** | Currant Creek | 39.8987 | -111.8891 | 0.5 | 0.00 | 1 | 0.00 |
| **CurFS-12** | Currant Creek | 39.8983 | -111.8888 | 0.5 | 38.03 | 1 | 59.11 |
| **CurFS-13** | Currant Creek | 39.8980 | -111.8884 | 0.5 | 0.00 | 2 | 0.00 |
| **CurFS-14** | Currant Creek | 39.8976 | -111.8881 | 0.5 | 0.00 | 0 | 0.00 |
| **CurFS-15** | Currant Creek | 39.8972 | -111.8878 | 0.5 | 0.00 | 0 | 0.00 |
| **CurFS-16** | Currant Creek | 39.8968 | -111.8878 | 0.5 | 39.31 | 1 | 40.77 |
| **CurFS-17** | Currant Creek | 39.8964 | -111.8882 | 0.5 | 39.83 | 1 | 29.48 |
| **CurFS-18** | Currant Creek | 39.8960 | -111.8884 | 0.5 | 42.36 | 0 | 2.02 |
| **CurFS-19** | Currant Creek | 39.8956 | -111.8886 | 0.5 | 0.00 | 0 | 0.00 |
| **CurFS-20** | Currant Creek | 39.8952 | -111.8888 | 0.5 | 0.00 | 0 | 0.00 |
